## Supplementary Material for "Targeting plasmid-encoded proteins that contain immunoglobulin-like domains to combat antimicrobial resistance"

**Table S1**. List of the strains and plasmids used in this work.

| Strain | Genotype and relevant properties | Reference |
|---|---|---|
| ***E. coli*** **DH10BT1R** | F- *mcrA* Δ*mrr-hsdRMS-mcrBC* ϕ80lacZDM15 Δ*lacX74* *recA1* *endA1* *araD139* Δ(*ara,leu*) *7697 galU galK rpsL (StrR) nupG tonA λ* | Invitrogen |
| ***E. coli*** **BL21DE3** | *hsdS, gal, (λclts857, ind1, Sam7, nin5, lac-UV5-T7gene1)* | (Studier and Moffatt, 1986) |
| ***S.*** **Typhimurium SL1344** | *rspLhisG* | (Hoiseth and Stocker, 1981) |
| ***S.*** **Typhimurium SL1344** ***ibpA::lacZ-Kmr*** | *rspLhisG ibpA::lacZ-Km$^r$* | (Hüttener et al., 2018) |
| ***S.*** **Typhimurium SL1344** ***flhDC::Km*** | *rspLhisG flhDC::Km* | This work |

| Plasmid | Description | Reference |
|---|---|---|
| **R27** | IncHI1, Tc$^r$ | (Grindley et al., 1972) |
| **R27 Δ*rsp*** | R27 *rsp*::FRT | (Hüttener et al., 2019) |
| **pHCM1** | IncHI1, Amp$^r$ | (Parkhill et al., 2001) |
| **pKD4** | *bla* (Ap$^r$) FRT *ahp* FRT PS1 PS2 oriR6K Km$^r$ | (Datsenko and Wanner, 2000) |
| **pKD46** | *bla* (Ap$^r$) $P_{BAD}$ *gam bet exo* pSC101 oriTS | (Datsenko and Wanner, 2000) |
| **pNeae2** | pNeae-derivative; for fusions to Neae-myc [$Intimin_{EHEC}$ (1-654)-E-His-myc tag | (Salema et al., 2013) |
| **pNeae2 VHH-RSP #3** | VHH against RSP protein fused to Neae-myc [$Intimin_{EHEC}$ (1-654)-E-His-myc tag | This work |
| **pNVFib1** | pNeae-myc-derivative; NVFib (clone 1) fusion [$Intimin_{EHEC}$ (1-654)-E-VFIBn-myc tag] | (Salema et al., 2016) |

| | | |
|---|---|---|
| **pMAL-RSP #5/7** | pMAL-p2E derivative, for fusion of C-terminal of RSP to MBP | This work |
| **pIgΔCH1** | pIgγ1HC derivative vector lacking human IgG CH1 domain | (Casasnovas et al., 2022) |
| **pIgΔCH1 VHH-RSP** | IgH signal peptide, VHH-RSP fused to the human IgG1 hinge and Fc portion (Fc) | This work |

**Table S2**. List of the oligonucleotides used in this work.

| Oligonucleotide | Sequence (5'-3') |
|---|---|
| **RSP5 BamHI Fw** | 5' CGGGATCCTGGGGTGTATAAGTTTCCT 3' |
| **RSP PstI Rv** | 5' AACTGCAGTTACTGCGAGGTTTCAAC 3' |
| **flhDC p1** | 5'GGCTACGTCGCACAAAAATAAAGTTGGTTATTCTGGATGGGAGTGTAGGC TGGAGCTGCTTC 3' |
| **flhDC p2** | 5'TTACCGCTGCTGGAGTGTTTGTCCACACCGTTTCGGTTAAACCATATGAAT ATCCTCCTTAGT 3' |
| **flhDC p1 up** | 5' CGTTGTATGTCACGAAGCTGAC 3' |
| **flhDC p2 down** | 5' GCTGTTGACTATGACAGGATGC 3' |
| **VHH Sfi** | 5' GTCCTCGCAACTGCGGCCCAGCCGGCCATGGCTCAGGTGCAGCTGGTG GA 3' |
| **VHH Not** | 5' GGACTAGTGCGGCCGCTGAGGAGACGGTGACCTGGGT 3' |
| **VHH pIg AgeI** | 5' ACTGCAACCGGTGTACATTCTCAGGTGCAGCTGGTGGA 3' |
| **VHH pIg BamHI** | 5' ACCGGATCCACGCGGAACCAGCGCTGAGGAGACGGTGACCTG 3' |
| ***Il-1* β Fw** | 5' GGTCAAAGGTTTGGAAGCAG 3' |
| ***Il-1* β Rv** | 5' TGTGAAATGCCACCTTTTGA 3' |
| ***Il*-6 Fw** | 5' TGTGAAATGCCACCTTTTGA 3' |
| ***Il*-6 Rv** | 5' GGTCAAAGGTTTGGAAGCAG 3' |
| *Tnf-α* Fw | 5' CCACCACGCTCTTCTGTCTAC 3' |
| *Tnf-α* Rv | 5' AGGGTCTGGGCCATAGAACT 3' |
| ***Hprt1* Fw** | 5' TGGATACAGGCCAGACTTTGTT 3' |
| ***Hprt1* Rv** | 5' CAGATTCAACTTGCGCTCATC 3' |

```
VHH-RSP#3      QVQLVESGGGFVQAGGSLRLSCAASGG--IG-DYRMAWFRQAPGKEREGVADINTSGGST
VHH-RSP#9      QVQLVESGGGSVQAGRSLRLSCAVSGY--SYSRRYMGFFRQAPGMEREGVAAISSTSGTT
VHH-RSP#20     QVQLVESGGGSVQVGGSLRLSCAASGD--FYRRDDVAWFRQAPGKEREGVAVIYVADGRT
VHH-RSP#21     QVQLVESGGGTVQIGGSLTLSCVGIGHNFWFSQLSLAWFRQAPGKEREGIAHITPSGQRT
VHH-RSP#22     QVQLVESGGGTVQVGGSLTLSCVGIGHNFWFSQLSLAWFRQAPGKEREGVARITPSGQGT
VHH-RSP#32     QVQLVESGGGSVQVGGSLRLSCAASGD--FYRRDDVAWFRQAPGKEREGVAVIYVADGRT
VHH-RSP#41     QVQLVESGGGSVQVGGSLRLSCAASGD--FYRRDDVAWFRQAPGKEREGVAVIYVADGRT
               ********** ** * ** ***.  *         :.:****** ****:* *  :.  *

VHH-RSP#3      RYVDSVKGRFTISRDNAKNTVYLQMNSLKPEDTGIYYCAAAPK-TQWARLQPESFNVWGQ
VHH-RSP#9      RYAESVKGRFIISRDTSKGTVYLQMNSLKLDDSAMYYCALARL-PSGNWLAASSYNYWGQ
VHH-RSP#20     RYVDSVKGRFTISYDSPKNTVYLQMNSLKSDDSAVYYCTMRRD-GGSRSWNPDRYTYWEQ
VHH-RSP#21     KYSETLEGRFTISRDNANNTMYLHMNNLKPEDAAMYYCAGAQSMGASQRLRAEKYAYWGQ
VHH-RSP#22     KYSETLEGRFTISRDNANSTMYLHMNNLKPEDAAMYYCAGGQSMGASQRLRAEKYSYWGQ
VHH-RSP#32     RYVDSVKGRFTISYDSPKNTVYLQMNSPKSDDSAVYYCTMRRD-GGSRSWNPDRYTYWGQ
VHH-RSP#41     RYVDSVKGRFTISYDSPKNTVYLQMNSLKSDDSAVYYCTMRRD-GGSRSWNPDRYTYWEQ
               :* ::::*** ** *. :.*:**:**. * :*:.:***:             . :  * *

VHH-RSP#3      GTQVTVSSAAAEQKLISEEDAAA    139
VHH-RSP#9      GTQVTVSSAAAEQKLISEEDAAA    140
VHH-RSP#20     GTQVTVSSAAAEQKLISEEDAAA    140
VHH-RSP#21     GTQVTVSSAAAEQKLISEEDAAA    143
VHH-RSP#22     GTQVTVSSAAAEQKLISEEDAAA    143
VHH-RSP#32     GTQVTVSSAAAEQKLISEEDAAA    140
VHH-RSP#41     GTQVTVSSAAAEQKLISEEDAAA    140
               ***********************
```

**Supplementary Figure 1. Amino acid sequence alignment of the selected VHHs after the MACS and FACS enrichment protocol of the bacterial library generated after the immunization of two dromedaries with the RSP protein.** Red color, blue color and purple color show the **CDR1**, **CDR2** and **CDR3** of the corresponding VHH, respectively. Yellow color marks the sequence of the c-myc tag used to evaluate the Nb display on the bacterial surface by flow cytometry. Alignment generated with Clustal Omega. Labels indicate full level of conservation (*) or decreased degree of conservation (: or .).

**Supplementary material references.**


Casasnovas JM, Margolles Y, Noriega MA, Guzmán M, Arranz R, Melero R, Casanova M, Corbera JA, Jiménez-de-Oya N, Gastaminza P, Garaigorta U, Saiz JC, Martín-Acebes MÁ, Fernández LÁ. 2022. Nanobodies Protecting From Lethal SARS-CoV-2 Infection Target Receptor Binding Epitopes Preserved in Virus Variants Other Than Omicron. *Front Immunol* **13**. doi:10.3389/fimmu.2022.863831

Datsenko KA, Wanner BL. 2000. One-step inactivation of chromosomal genes in Escherichia coli K-12 using PCR products. *Proc Natl Acad Sci U S A* **97**:6640–6645. doi:10.1073/pnas.120163297

Grindley NDF, Grindley JN, Anderson ES. 1972. R factor compatibility groups. *MGG Molecular & General Genetics* **119**:287–297. doi:10.1007/BF00272087

Hoiseth SK, Stocker BAD. 1981. Aromatic-dependent Salmonella typhimurium are non-virulent and effective as live vaccines. *Nature* **291**:238–239. doi:10.1038/291238a0

Hüttener M, Prieto A, Aznar S, Bernabeu M, Glaría E, Valledor AF, Paytubi S, Merino S, Tomás J, Juárez A. 2019. Expression of a novel class of bacterial Ig-like proteins is required for IncHI plasmid conjugation. *PLoS Genet* **15**:e1008399. doi:10.1371/journal.pgen.1008399

Hüttener M, Prieto A, Aznar S, Dietrich M, Paytubi S, Juárez A. 2018. Tetracycline alters gene expression in Salmonella strains that harbor the Tn10 transposon. *Environ Microbiol Rep* **10**:202–209. doi:10.1111/1758-2229.12621

Parkhill J, Dougan G, James KD, Thomson NR, Pickard D, Wain J, Churcher C, Mungall KL, Bentley SD, Holden MTG, Sebaihia M, Baker S, Basham D, Brooks K, Chillingworth T, Connerton P, Cronin A, Davis P, Davies RM, Dowd L, White N, Farrar J, Feltwell T, Hamlin N, Haque A, Hien TT, Holroyd S, Jagels K, Krogh A, Larsen TS, Leather S, Moule S, Ó'Gaora P, Parry C, Quail M, Rutherford K, Simmonds M, Skelton J, Stevens K, Whitehead S, Barrell BG. 2001. Complete genome sequence of a multiple drug resistant Salmonella enterica serovar Typhi CT18. *Nature* **413**:848–852. doi:10.1038/35101607

Salema V, López-Guajardo A, Gutierrez C, Mencía M, Fernández LÁ. 2016. Characterization of nanobodies binding human fibrinogen selected by E. coli display. *J Biotechnol* **234**:58–65. doi:10.1016/j.jbiotec.2016.07.025

Salema V, Marín E, Martínez-Arteaga R, Ruano-Gallego D, Fraile S, Margolles Y, Teira X, Gutierrez C, Bodelón G, Fernández LÁ. 2013. Selection of Single Domain Antibodies from Immune Libraries Displayed on the Surface of E. coli Cells with Two β-Domains of Opposite Topologies. *PLoS One* **8**:e75126. doi:10.1371/journal.pone.0075126

Studier FW, Moffatt BA. 1986. Use of bacteriophage T7 RNA polymerase to direct selective high-level expression of cloned genes. *J Mol Biol* **189**:113–130. doi:10.1016/0022-2836(86)90385-2
